# GPFN: Prior-Data Fitted Networks for Genomic Prediction

**DOI:** 10.1101/2023.09.20.558648

**Authors:** Jordan Ubbens, Ian Stavness, Andrew G. Sharpe

## Abstract

Genomic Prediction (GP) methods predict the breeding value of unphenotyped individuals in order to select parental candidates in breeding populations. Among models for GP, classical linear models have remained consistently popular, while more complex nonlinear methods such as deep neural networks have shown comparable accuracy at best. In this work, we propose the Genomic Prior-Data Fitted Network (GPFN) as a new paradigm for GP. GPFNs perform amortized Bayesian inference by simulating hundreds of thousands or millions of plant or animal populations. This allows GPFNs to be deployed without requiring any training or tuning, providing predictions in a single inference pass. On three populations of plants across two different crop species, GPFNs perform significantly better than the linear baseline on 13 out of 16 traits. On a challenging between-families structured prediction task on a third crop species, the GPFN matches the performance of the linear baseline while outperforming it in one location. GPFNs represent a completely new direction for the field of genomic prediction, and have the potential to unlock levels of selection accuracy not possible with existing methods, especially in diverse populations.

## 1. Introduction

With the goal of supporting a growing global population, increasing the rate of genetic gain in plant and animal breeding is important. In modern breeding, Genomic Selection (GS) has become one point of focus for improving genetic gain in quantitative traits [1]. GS aims to select the most fit parents by performing Genomic Prediction (GP) – regressing phenotypic values using genome-wide marker data, most often Single-Nucleotide Polymorphisms (SNPs). The efficacy of GS has been validated in several plant and animal species, accelerating the rate of genetic gain in many cases [2]–[4]. With genotyping becoming increasingly accessible through technological improvement and cost reduction, GS has become relevant for more and more breeding programs in recent years.

Despite continual progress in genotyping, the prediction of genotypic value is still most often performed using classical techniques such as BayesB [1], and GBLUP [5]. Much recent work has focused on the discovery of new methods based on deep learning [6]–[8] – however, classical methods remain the most popular because they empirically perform at least as well as newer machine learning-based methods while being simpler, faster, and requiring less tuning [9]–[14].

In all cases in the current literature, GP is carried out in a supervised fashion, where a model is trained using the training population and then that model is used to perform inference on the target (testing, evaluation) population. The learning behavior of any particular estimator is influenced by the model’s priors, such as inductive biases expressed through model architecture or regularization schemes. These priors are usually especially important when it comes to high-capacity models such as deep neural networks operating in low-data regimes. The alternative to these types of implicit priors is Bayesian inference, where the prior belief over the data is defined explicitly. This is straightforward if the data generating process can be modelled as a tractable distribution, such as a mixture of Gaussians, but for more complex distributions (such as those found in quantitative genetics) it is difficult to estimate the Posterior Predictive Distribution (PPD).

One way to overcome this limitation is with the recently described *Prior-Data Fitted Networks* (PFNs), which allow a neural network to approximate the PPD given any prior from which one can sample [15]. The prior can be intractable, such as a complex, black-box process, and the PFN is still able to perform approximate Bayesian inference. Practically, the process of training the PFN involves repeatedly sampling datasets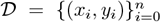, as input-label pairs, from the prior distribution over datasets *p*(). During training, the *n* datapoints are used for training along with *query points x* for which we wish to estimate the PPD *p*(*y* | *x*, 𝒟). The PPD is given by

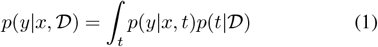

where *p*(*t*) is a prior for a latent variable, called the task [15]. One can think of the task as a particular internal state of the data-generating process.

PFNs model the PPD, without ever instantiating the posterior, parameterized by a neural network. In particular, PFNs use the transformer architecture [16] as it is well-suited to modeling relationships between the set-ordered datapoints in the datasets drawn from the prior. During training, datasets 𝒟 are sampled from the prior *p*(𝒟) and model parameters *θ* are optimized by minimizing

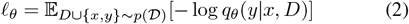

via gradient descent. It is shown that optimizing Equation 2 is equivalent to minimizing the cross-entropy between the PPD and its approximation [15].

In this work, we develop a new approach to GP based on amortized Bayesian inference, which we term *Genomic Prior-Data Fitted Networks* (GPFNs). Unlike existing methods based on neural networks, a GPFN is not trained on the end user’s dataset, does not require any tuning, and in fact is never exposed to any real data at all prior to inference. The key difference between the classical supervised learning approach typically used in GP and the Bayesian inference performed by PFNs is that beliefs about factors such as population structure and recombination are expressed implicitly via the prior. As a proof of concept, we show that the GPFN is the first method which is able to consistently and significantly outperform classical methods in several datasets of crop plants. We propose the GPFN approach as a major new direction for genomic prediction. A full implementation as well as several trained GPFNs for various population types are provided at https://github.com/jubbens/gpfn.

## II. Materials and Methods

### PFNs for Genomic Prediction

GPFNs are fit using priors which are defined by different types of simulated populations. As a convention, we use the term *unstructured population* to refer to a population which is not the result of a defined crossing scheme, such as diversity panels, and *structured population* to refer to those which are, such as genetic mapping panels and recurrent breeding populations. Because different populations vary in a variety of ways, a prior must be created specifically for each type of population. When creating the prior, the samples must be split with the same training and testing split as will be seen during inference. For example, if one were to train a GPFN for use in a breeding program where the F4s from the previous recurrent block are used as a training population to perform selection in the F2s in the subsequent recurrent block, the prior should ideally be designed to simulate this exact procedure. In the next section we discuss two priors built for demonstrating GPFNs: one for unstructured populations with random train/test splits (which we will demonstrate in three unstructured populations of wheat and lentil) and one for structured NAM populations with a between-families train/test split (which we will demonstrate in a soybean NAM population).

### Priors for Prediction Populations

PFNs have been fit using priors defined by Bayesian neural networks, Gaussian processes, and structural-causal models [15], [17]. In order to create priors which model plant and animal populations, we use genetic simulations based on SeqBreed [18]. In the context of GPFNs, a prior is defined by a simulator which simulates a particular type of population (wild, mapping, breeding, etc.) and each draw from the prior provides a completely new population of this type. Each prior has a number of parameters which define characteristics of the quantitative trait architecture, population structure, and other factors (Table I). When drawing from the prior, these parameters are sampled from a uniform distribution.

For simplicity, all priors demonstrated here assume that experiments are single-trait, single-environment, and unreplicated. Multi-environment and multi-trait models are left for future work. The quantitative trait is simulated through SeqBreed. To define the quantitative trait architecture, we simulate both additive and dominance effects (with the proportion of dominant alleles being a randomly sampled parameter). Marker effects are sampled from a gamma distribution [18], with the *α* and *β* parameters being randomly sampled. The trait heritability (*h*^2^) and the number of quantitative trait nucleotides (QTN) are also randomly sampled. The diploid genome of the population is also simulated through SeqBreed, which also handles all of the simulated crossing and recombination, with the total number of SNPs, the recombination rate, the heterozygosity, and the minor allele frequency all being randomly sampled parameters.

The final outputs for a single draw from the prior are the SNP markers and phenotypic values for the members of the training and testing populations. In the following sections we discuss the details for how priors are defined for two different types of populations: unstructured (wild) populations, and nested association mapping (NAM) populations.

### Unstructured Population Prior

To train a GPFN for use with unstructured populations, we use a prior which simulates a wild population. All priors for structured populations (such as the NAM prior) also start with founders drawn from this prior (see Figure 1). Although this is not a realistic starting point for breeding populations which typically start from elite inbred lines, it is more difficult to generate these types of parents *de novo*.

**Fig. 1.**
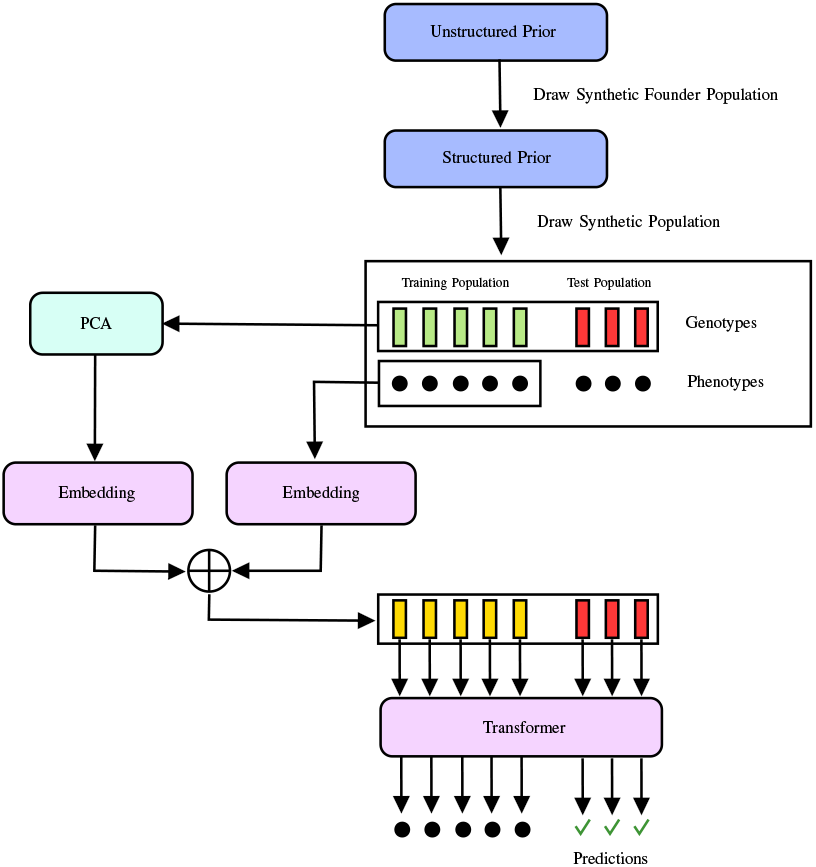
Training a Prior-Data Fitted Network to perform Genomic Prediction. A single draw from the prior is shown here for illustrative purposes; during fitting, this process is repeated one million times and multiple draws are organized into training batches. The founders for populations from the structured prior are drawn from an unstructured prior. For unstructured populations, the unstructured prior is used directly. We use linear projections for the embedding modules. The transformer model takes the training and query individuals, with the training individuals conditioned by their phenotype label by adding the label embedding. Finally, the transformer outputs phenotype predictions for the query individuals (red).

In order to generate the unstructured population, we begin from a random number of individuals with randomized biallelic SNPs. For a random number of generations, we proceed with random mating. Each cross between these random pairings results in between 1 and 4 children. With a randomly sampled probability, half of the current population migrates to found a new simulated subpopulation. This process proceeds for each time step, with parents being randomly selected within each distinct subpopulation. Allele frequency begins to differentiate between the different subpopulations due to reproductive isolation. Finally, extant individuals are randomly selected across the different subpopulations and randomly apportioned to the training and test sets. Pseudocode for the complete process is shown in Algorithm 1. The random parameters for the unstructured prior are listed in Table I. Figure S1 shows an example of a population drawn from the unstructured prior compared to a real population of wheat landraces.

### NAM Prior

The Nested Association Mapping prior models a standard NAM scheme, where prediction is performed between-families. First, founders are generated by drawing from the unstructured prior. These individuals are not split into training and testing, as this split will happen on the families of the future NAM population. Of the founders, one is selected at random as a common parent and the others are crossed to it, followed by single seed descent. The total number of families is randomly varied between 25 and 85, while the number of iterations of selfing is randomly varied between 2 and 5. The extant individuals are grouped by family, and we split the families into training and testing. All other parameters are identical to those in the unstructured prior (Table I). Figure S2 shows an example of a population drawn from the NAM prior compared to a real NAM population of soybean.

### Prior Fitting

To fit a GPFN model we largely follow the fitting procedure from Müller et al [15]. During model fitting, the prior is repeatedly sampled to generate new 𝒟 which constitute training batches. The GPFNs presented here are fit using one million draws from the prior. Figure 1 illustrates an overview of the prior fitting process.

For the neural network component, PFNs use transformer models [16] to approximate the PPD. The use of transformers as PFNs comes with some practical computational challenges. Since the memory complexity of self-attention scales quadratically with the number of tokens, the number of individuals *n* per sampled is 𝒟 limited. Because the input tokens are set-valued, *i*.*e*. not ordered, it is not viable to use faster approximations which focus on local attention windows. We limit the population size to a maximum of 1000 during training, but during inference we use the full training and testing sets. The size of the input vectors *x*_*i*_ needs to be constant despite the different number of SNPs per simulated population. To achieve this, we use the loadings on the first 100 principal components as the input to the embedding layer.

The number of query points is varied between 10 and 500 during training, although we use a uniform weighting over the number of training versus query samples instead of one biased towards larger training sets as in Müller et al [15]. This is because we prefer that the network is frequently exposed to smaller training populations due to the interaction between the number of samples in the training set and the PCA projection step.

PFNs are usually trained using the Prior-Data Negative Log Likelihood loss (Equation 2). However, we find that using a bucketized mean squared error provides slightly better performance on our regression task. This involves computing the mean prediction by weighting the bucket centers by the output probability, instead of only considering the probability attributed to the correct bucket. Although it breaks some theoretical guarantees, we find that this stabilizes training as well as allowing for training under a wider range of hyperparameters, since the gradient is smoother under the bucketized MSE criteria.

### Architectural Details

We fit the prior using an encoder-only transformer model with 12 layers, an internal dimensionality of 2048, a hidden layer size of 2048, and a single attention head. In our experiments, multi-head attention did not yield better results than a single attention head, so a single head is used for faster throughput. In general, wider and deeper transformers performed slightly better when using the same prior. This is despite the fact that the smaller transformers with fewer parameters were usually able to fit the prior to a similar level of accuracy during the fitting phase. This is why we use an architecture which is both wider and deeper than PFNs demonstrated in the literature [15].

### Evaluation Strategy

For all results, we compare to GBLUP, because we consider it to be at least as good as state-of-the-art. We also compare to Principal Component Regression (PCR), since the GPFN uses the PCA loadings as input, as this helps elucidate how much of the performance difference may be attributable to the projection method instead of the inference method. Additionally, PCR shows what level of performance can be attained by simply performing linear regression on these features. This helps to show whether or not the transformer is implementing a prediction strategy which is superior to linear regression. We also compare to XGBoost [19] as a machine learning baseline. Since XGBoost ideally requires hyperparamter tuning, we use a 3-fold tuning strategy with random search and a budget of 200 iterations. We search over the number of estimators, learning rate, maximum depth, and the data subsample ratio. Following tuning, we apply the model with the best cross-validation performance to the test set. It is not our intention to compare across all classes of models, as this has already been demonstrated extensively in the literature [9]–[14]. For each experiment, we split the data into 80% training and 20% testing. We measure the Pearson’s correlation between the estimated genotypic value output by the model and the observed phenotypic value. The data is resampled 100 times to obtain error estimates. Full results are provided in Supplementary Note C.

## III. Results

### Wheat Preliminary Yield Trials

We examined performance on a breeding population using a preliminary yield trial (PYT) of wheat from the University of Guelph’s Winter Wheat Breeding program [20]. The PYT is a single-rep, single-environment trial. In total, this population is comprised of 319 individuals and 4,722 SNPs. The population is sourced from diverse sources, and the authors infer a low level of relatedness in the PYT population due to a small amount of variance explained by principal components 1 and 2. For all trials including the PYT, broad-sense heritability (*H*^2^) of the traits under consideration varied between 0.13 for yield, 0.97 for thousand kernel weight, 0.87 for maturity, and 0.98 for plant height. We use a GPFN fit with the unstructured prior for this evaluation. Figure 2 shows results for the wheat PYT population.

**Fig. 2.**
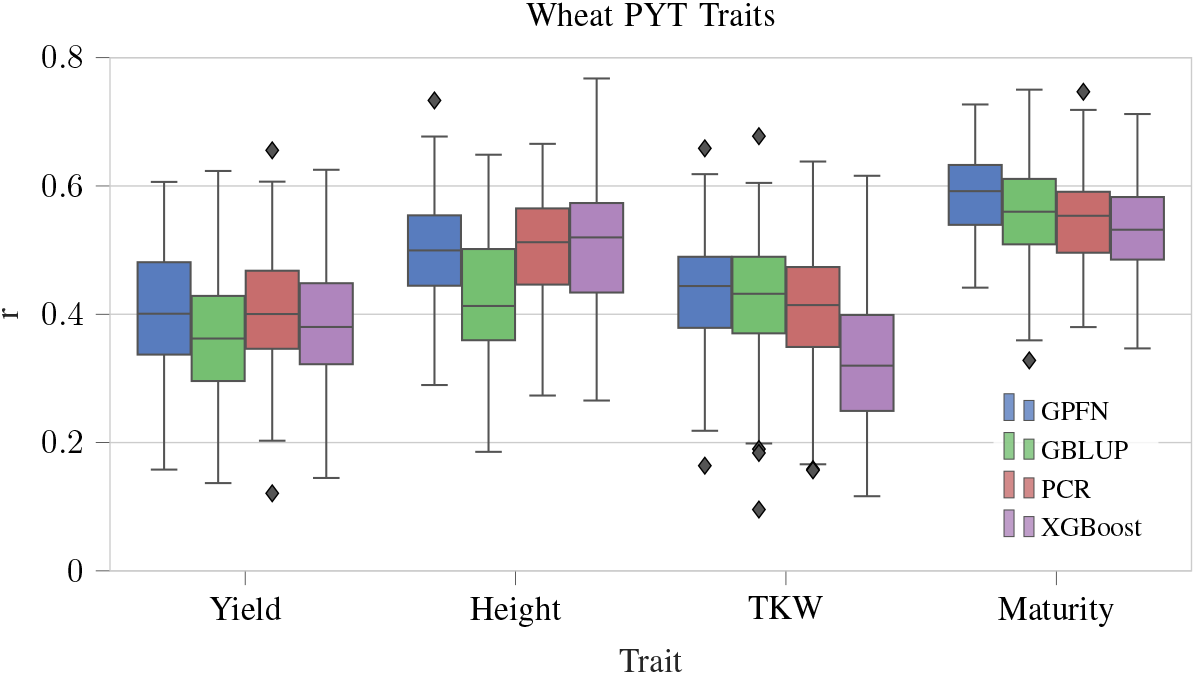
Prediction performance results for the wheat preliminary yield trial. Detailed results are shown in Table II.

### Wheat Landraces

For another unstructured crop population, we use a dataset containing 2,000 Iranian landrace accessions of wheat from the CIMMYT gene bank [21]. The authors note the presence of population structure in the landraces. We selected the six traits which were measured in only a single environment. Narrow sense heritabiliy (*h*^2^) was 0.839 for grain hardness, 0.881 for grain length, 0.434 for plant height, 0.625 for grain protein, 0.754 for test weight, 0.833 for thousand-kernel weight, and 0.848 for grain width. The data consists of 33,709 SNPs. We use a GPFN fit with the unstructured prior for this evaluation. Figure 3 shows results for the wheat landraces population.

**Fig. 3.**
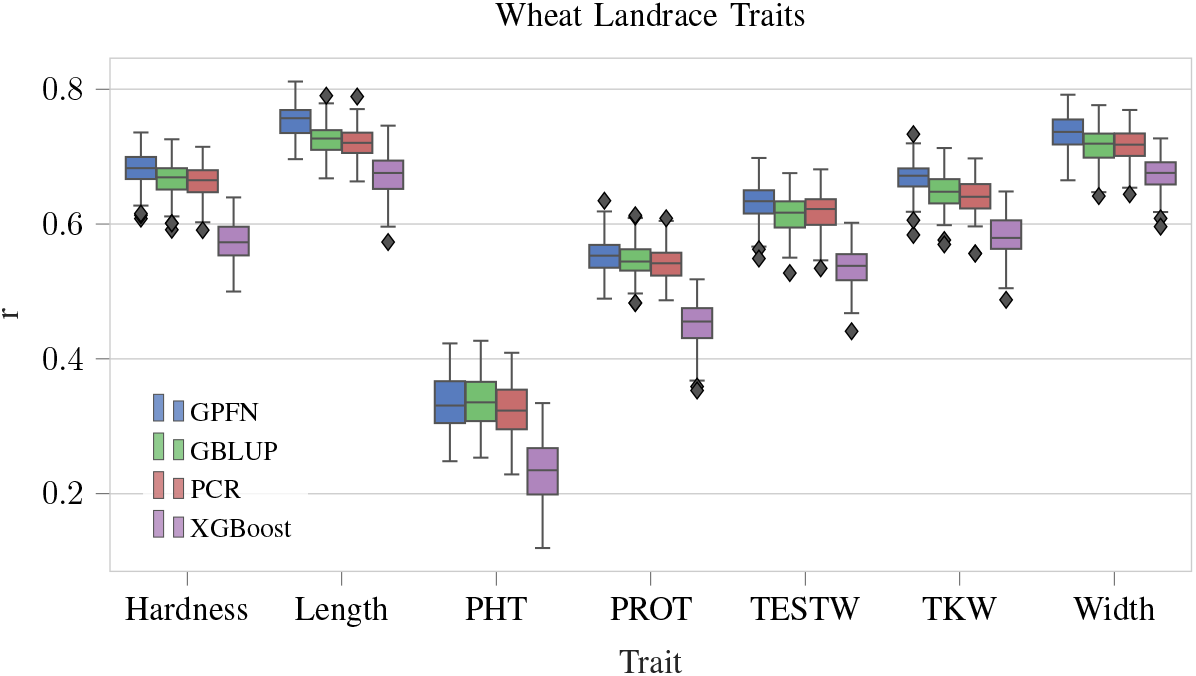
Prediction performance results for the wheat landraces. Detailed results are shown in Table III.

### Lentil Diversity Panel

For the final unstructured crop population, we use a diversity panel of lentil [22]. This is a very diverse population, assembled from various global gene banks as well as select breeding lines from a public lentil breeding program in Canada. The authors use multiple environments to estimate the broad-sense heritability (*H*^2^) of the measured traits to be 0.81 for days to flowering (DTF), 0.82 for vegetative period (VEG), 0.74 for days to swollen pods (DTS), 0.62 for days to maturity (DTM), and 0.29 for reproductive period (REP). Since we are interested in the single-environment case, we select the Rosthern 2017 environment as it has measurements available for all traits. There are a total of 324 individuals and 9,394 SNPs. We use a GPFN fit with the unstructured prior for this evaluation. Figure 4 shows performance results for the lentil diversity panel.

**Fig. 4.**
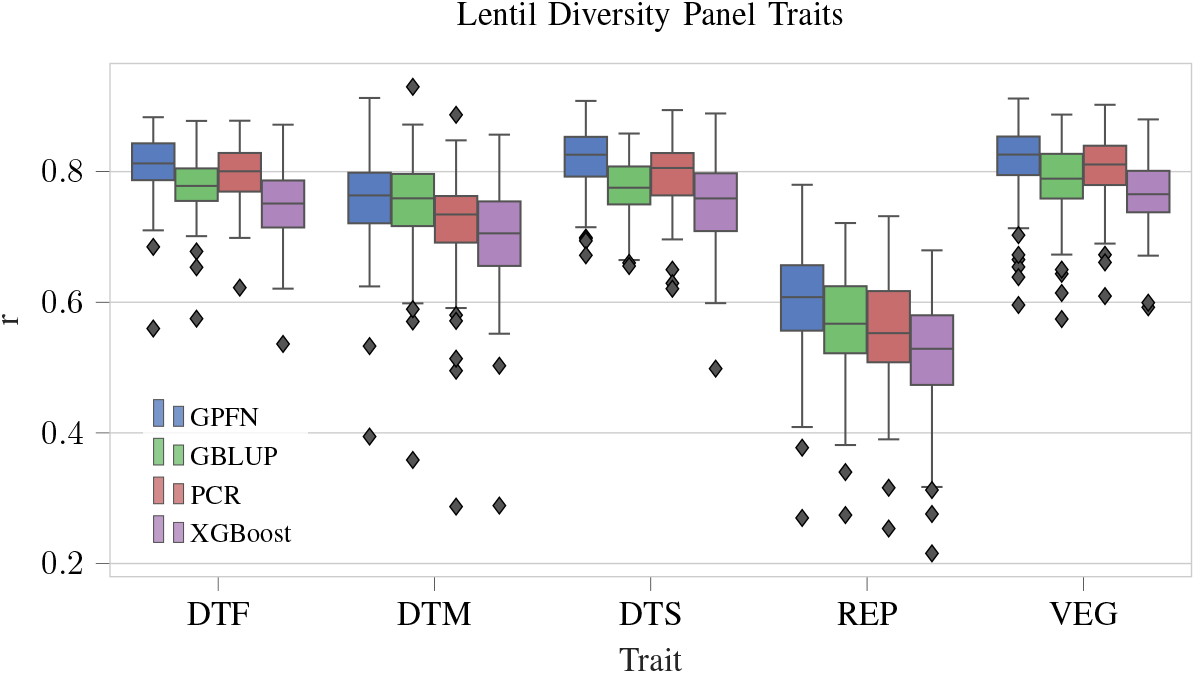
Prediction performance results for the lentil diversity panel. Detailed results are shown in Table IV.

### Nested Association Mapping Panel of Soybean

For a structured population, we use a nested association mapping (NAM) panel of soybean, known as SoyNAM [23]. Because the published marker data contains a high proportion of missing data, we use a version with imputed genotypes [24]. For the trait of interest, we predict yield across multiple locations. We use the 2012 year since data is available for all locations for this field season. This population contains a variable number of individuals between locations, from 827 in the MO location to 5019 in NE. The total number of families also varies between locations, with 23 in MI, 25 in MO, and 38 in each of OH, IL, IA, IN, NE, and KS. The imputed genotype data consists of 29,416 SNPS.

This population has been used extensively for genomic prediction in past studies [14], [25]. We use a between-families prediction scheme as this represents one of the most difficult prediction tasks, owing to the difficulty of predicting between families, as well as using a trait which is only moderately heritable and shows low prediction accuracy in general. This is similar to the leave-family-out sampling previously described in [25], but even more difficult as 20% of the families are held out for the evaluation population. We use a GPFN fit with the NAM prior for this evaluation. Figure 5 shows a summary of prediction results for the NAM population.

**Fig. 5.**
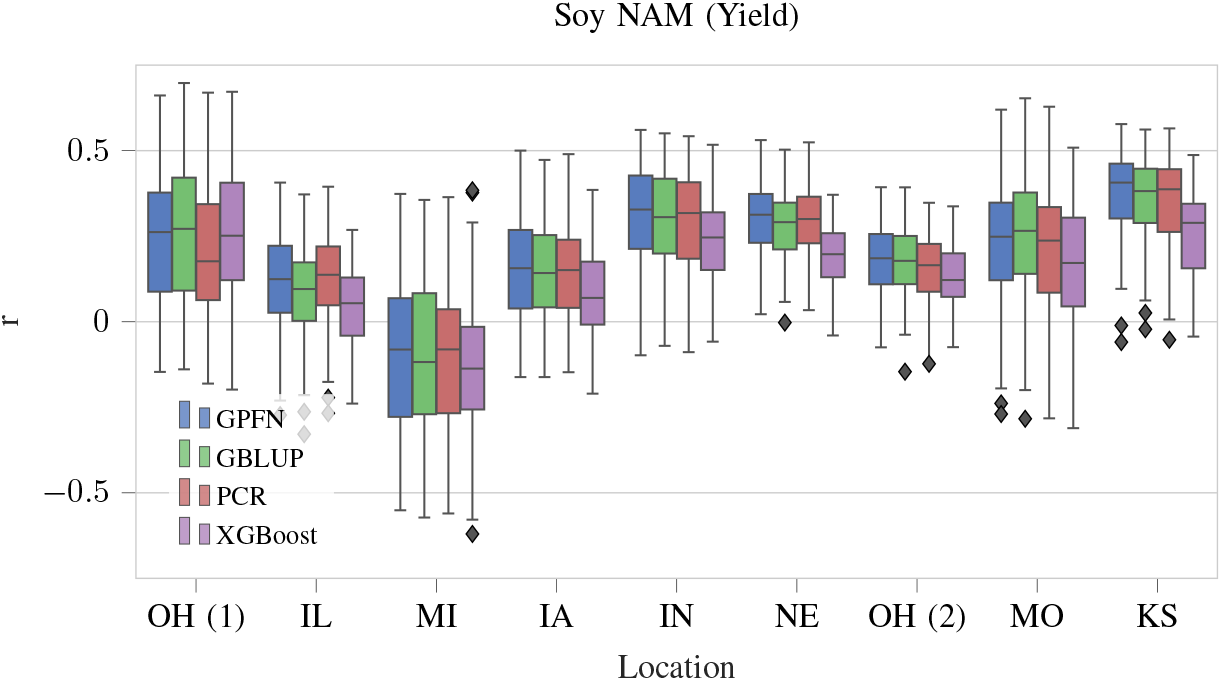
Prediction performance results for the soybean NAM population. Detailed results are shown in Table V. **OH**: Ohio; **IL**: Illinois; **MI**: Michigan; **IA**: Iowa; **IN**: Indiana; **NE**: Nebraska; **MO**: Missouri; **KS**: Kansas.

## IV. Discussion

### Performance Versus Baselines

For the GPFN fit using the unstructured prior and applied to the wheat PYT, wheat landraces, and lentil diversity panel datasets, the proposed method significantly outperformed GBLUP in 13 out of 16 traits, and significantly underperformed in none (Supplementary Note C). It is interesting to note that the GPFN trained on the unstructured prior shows generalization performance outside of the range of populations represented by its prior. For example, for the small wheat PYT and lentil diversity panel datasets, the size of the training set is significantly smaller than the smallest datasets the GPFN was exposed to during training. Similarly, for the wheat landraces dataset, the training set is significantly larger. The PYT data also contains far fewer SNPs than were simulated. Perhaps the most relevant difference is the types of individuals – the unstructured prior simulates a wild population with random mating, while the individuals in the real datasets are likely far more inbred. Despite these mismatches between the real data and the synthetic populations which the GPFN encountered during prior fitting, the GPFN exceeds the baseline performance convincingly in every case. There may be more performance available if these types of populations can be simulated more accurately.

For the GPFN fit using the NAM prior and applied to the SoyNAM dataset, the results were more mixed. The GPFN significantly outperformed GBLUP in predicting yield at only one of the nine locations, although it did not significantly underperform GBLUP in any. The remaining eight locations showed no statistically significant difference. Compared to the wheat PYT, wheat landraces, and lentil diversity panel which were tested using a GPFN fit with the unstructured prior, the GPFN fit with the NAM prior shows less of an advantage over GBLUP in the soybean NAM population. We believe that this is not necessarily due to a distinct weakness of the GPFN, but rather a distinct strength of GBLUP in these types of structured populations. It may be that, owing to the presence of a common parent, there exists a strong correlation between the linear covariance in the marker data and the underlying genotypic value. This relationship may be more complex in unstructured populations, where the GPFN tends to pull ahead.

The GPFN performs better than PCR across all datasets and traits with the exception of three, implying that the performance differences between GPFN and GBLUP are not attributable to PCA and that the model is implementing an algorithm which is superior to linear regression. XGBoost consistently underperformed both the GPFN and GBLUP across the different datasets, outperforming GBLUP in only one trait and underperforming it in a further 21.

### GPFNs Shift the Focus from Prediction Models to Simulation

Much effort has been dedicated to developing models which are well-suited to genomic prediction. This includes linear models which imply different distributions of marker effects (GBLUP, LASSO, BayesA, BayesB), as well as neural network methods which attempt to propose the “right” cocktail of regularization and architectural modifications to capture important information without overfitting the training set [6]– [8]. However, such an iterative approach where a model is tweaked until performance is improved on a particular dataset is more likely to result in a model which is tailored to a particular dataset or the specific trait architectures therein than it is to result in a model with good generalization capabilities. In the worst case, such an approach samples mostly noise, where the individual changes have effectively zero effect size, but eventually a model which performs better than the competing models on the target dataset is found by chance.

The GPFN approach is fundamentally different: the model architecture is held constant, while the prior is made as broad and realistic as possible. Overfitting is no longer an issue, as training data is practically infinite (as the prior can be sampled any number of times) and PFNs are intended to fit the prior as closely as possible [15]. The more diverse the prior is, the wider the range of quantitative trait architectures it is expected to generalize to. The proposed approach effectively shifts the focus from blindly searching over model architectures to creating the most accurate possible simulation models for quantitative traits.

### Additional Evidence Against the Importance of Marker Effects

Evidence has been presented that, when tasked with genomic prediction, deep neural networks tend not to estimate marker effects [12]. Despite being indifferent to marker effects, these networks still perform comparably to classical linear methods. Similarly, GPFNs are also unable to estimate the effects of individual markers due to the PCA projection. We show that they can perform even better than existing methods by imposing prior beliefs over populations instead of marker effects. This is corroborating evidence that, for polygenic traits, marker effects are less relevant for prediction than the broad patterns of relatedness implied by the markers.

### GPFNs that Fit the Prior Better Also Perform Better on Real Populations

The performance of a PFN should increase with the number of draws from the prior, as it fits the prior more tightly. Using the wheat landraces dataset, we tested the generalization ability of a GPFN at different points during prior fitting (Figure 6). It is clear that generalization does improve over the course of prior fitting, with more performance potentially available with additional fitting beyond one million draws.

**Fig. 6.**
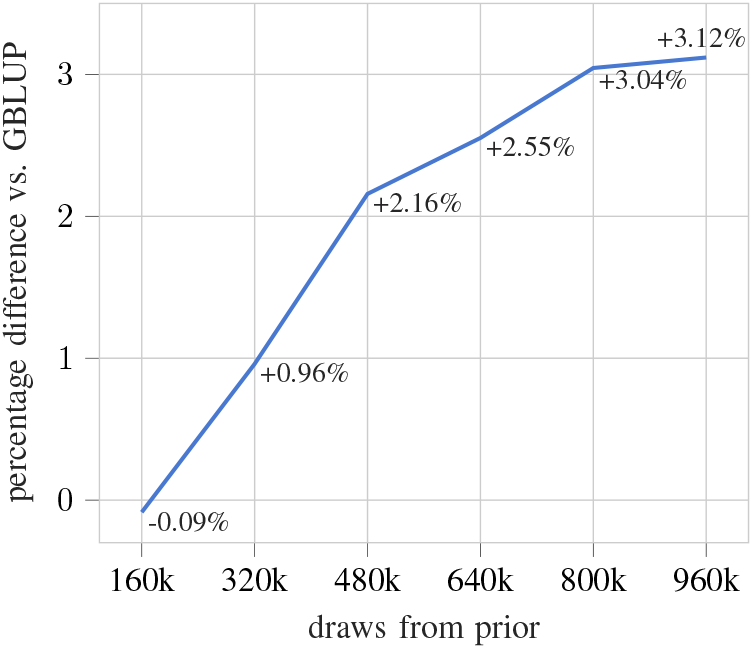
Performance on the wheat landraces dataset with a 50-50 train-test split, as a function of draws from the prior. Mean performance difference across all traits is shown.

### Training Time and Resources

Simulation was performed using 336 CPU cores, while fitting was done in parallel on a single V100 GPU. Using this configuration, the two models demonstrated here each required about seven days to generate one million draws from the respective prior. Although the model fitting process is relatively computationally intensive, it is important to remember that this cost is amortized: once the model is fit, its weights are saved and it can be reused indefinitely on any relevant populations without needing to be re-trained.

The benefits of this amortization were seen during our own evaluations, which required resampling each population 100 times for each trait, resulting in a total of 2,500 evaluations. Because GPFNs only perform inference, and do not need to be re-trained or tuned, each evaluation took only a few seconds to complete. On the larger populations, such as the wheat landraces, this was in fact even faster than the linear models (GBLUP and PCR). Other machine learning methods not only need to be re-trained each time, but a hyperparameter search also has to be performed independently on each random split of the data to avoid leaking information across training and testing sets.

## Limitations

The GPFN approach comes with some novel and practical limitations. Using PCA projection before embedding means that the size of the training set must be at least as large as the number of components used. For example, when 100 principal components are used, the training set must be comprised of at least 100 individuals. This is not a limitation shared by competing methods such as GBLUP, which do not perform dimensionality reduction prior to model fitting.

As with any simulation study, the relevance of the prior (and therefore the expected performance of the model) is dependent on the accuracy of the simulation. We endeavoured to represent a wide range of trait architectures in our simulations, but it is impossible to capture the full range of diversity present in real-world scenarios. For example, as our simulator is focused on quantitative traits, performance may suffer when presented with discrete or ordinal phenotypes (such as measurements in days, or disease ratings) as these types of phenotypes were not simulated during model fitting.

As mentioned previously, the prior needs to match the population well or performance will be degraded. For example, using a GPFN trained on the unstructured prior to perform inference on the between-families NAM structured population resulted in a mean performance shortfall of almost 15% versus the baseline (data not shown). For this reason, a prior should ideally be made for each unique breeding scheme in order to maximize the performance of the technique. Some standard breeding schemes can be modelled and reused for those applications – however, if the breeding scheme is nonstandard or the training and testing individuals are not split between different subpopulations in the same way, a new prior needs to be created. Another implication is that, in some breeding situations, the prior will need to be updated and the model refit on a yearly basis as new crosses are made with the current progeny. Otherwise, structure of the prior will drifter further and further from the reality of the breeding population with each year.

Although this initial proof of concept has demonstrated encouraging results with two priors, more evaluations are required on more types of structured and unstructured populations. In particular, more evaluations are required for breeding populations, such as those found in recurrent breeding programs. The lack of this type of data in the public domain means that the community will need to pursue these evaluations to further validate the GPFN approach in their own populations. We anticipate that the ability of GPFNs to model the idiosyncrasies of these structured populations will unlock even more prediction performance and genetic gain in the future.

## Acknowledgments

We would like to acknowledge Alexandra Ficht for providing the data for the wheat preliminary yield trial.

