## Supplementary Materials for "GPFN: Prior-Data Fitted Networks for Genomic Prediction"

### Appendix A
### Supplementary Figures

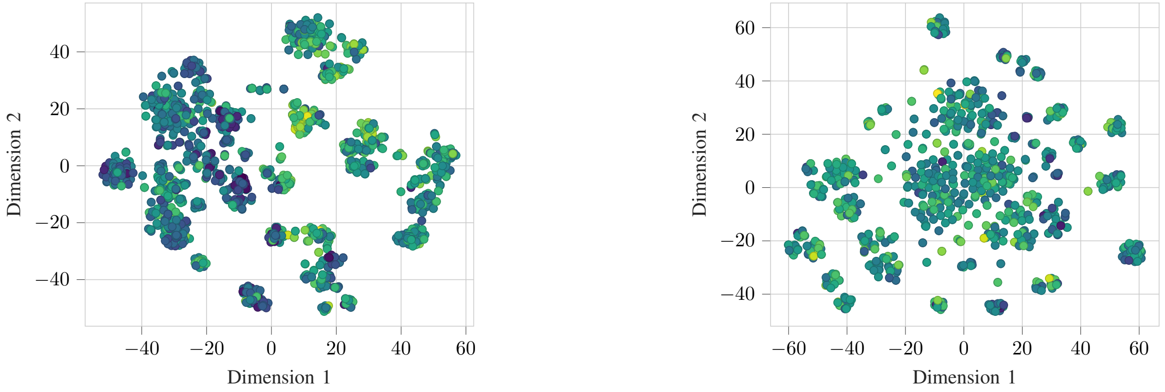


Fig. S1. The wheat landrace population (left) and a draw from the unstructured prior (right) visualized with t-SNE. Points are shaded by phenotype (grain hardness for the real population).

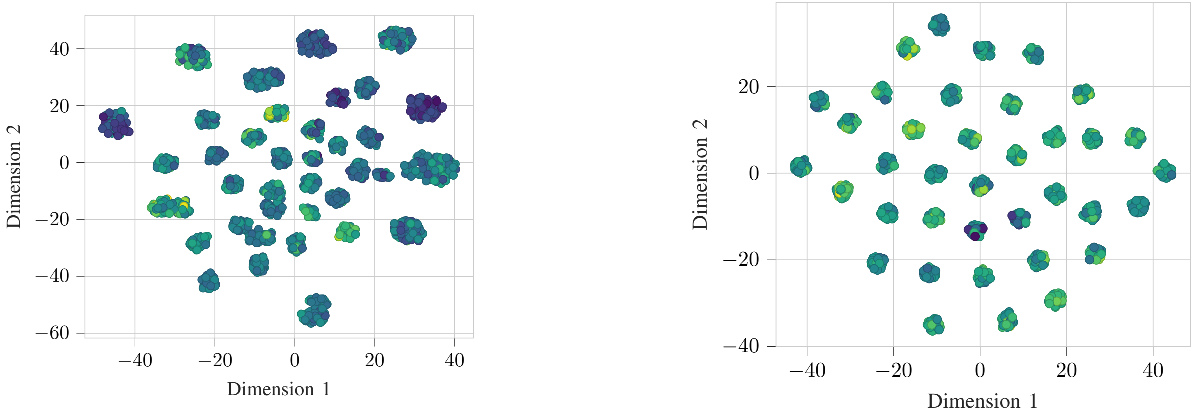


Fig. S2. The soybean NAM population (left) and a draw from the NAM prior (right) visualized with t-SNE. Points are shaded by phenotype (yield for the real population).

#### APPENDIX B
#### SIMULATION DETAILS

**Algorithm 1:** Drawing a sample from the wild prior. Parameters are listed in Table I. All genetic and quantitative trait simulation is done through SeqBreed.

```
Result: Markers and phenotypic values for training and testing populations
draw random values for all parameters from uniform distributions;
define a genome with H, MAF, numsnps, recomb.rate;
define a quantitative trait with h^2, numqtl, pdom, alpha, beta;
generations ← empty tree;
subpopulations ← empty tree;
founders ← population of size numbase with random alelles at all sites;
INSERT(generations, founders);
INSERT(subpopulations, founders);
for timesteps do
    for subpopulation in subpopulationsgraph do
        generation ← youngest generation in generationsgraph;
        select random parents from generation in subpopulation;
        progeny ← 1 to 4 children from each cross between parents;
        if random > psplit then
            split half of progeny into new branch;
            INSERT(subpopulations, branch);
        end
    end
    INSERT(generations, all progeny in all subpopulations);
end
generation ← youngest generation in generations;
extant ← all individuals in generation for each supopulation in subpopulations;
training ← random selection from extant;
testing ← random selection from extant;
```

| **timesteps** | **p split** | **num base** | **num snps** |
|---|---|---|---|
| [20, 60] | [0.1, 0.3] | [500, 1000] | [7000, 36000] |
| $h^2$ | $H$ | **p dom** | **alpha** |
| [0.2, 0.8] | [0.05, 0.15] | [0.0, 0.25] | [0.1, 0.3] |
| **beta** | **recomb. rate** | **num qtl** | **MAF** |
| [1.0, 9.5] | [0.5, 2.5] | [5, 100] | [0.1, 0.3] |

TABLE I
PARAMETERS FOR PRIORS. ALL PARAMETERS ARE SAMPLED UNIFORMLY. **P SPLIT**: PROBABILITY OF SPLITTING INTO A NEW SUBPOPULATION; **NUM BASE**: NUMBER OF INDIVIDUALS IN THE FOUNDING POPULATION; **P DOM**: PROBABILITY OF DOMINANCE EFFECT; **ALPHA, BETA**: PARAMETERS OF GAMMA DISTRIBUTION FOR MARKER EFFECTS; **RECOMB. RATE**: RECOMBINATION RATE IN CM/MBP; $H$: HETEROZYGOSITY; **MAF**: MINOR ALLELE FREQUENCY.

#### APPENDIX C
#### DETAILED RESULTS

| | GPFN | GBLUP | PCR | XGBoost |
|---|---|---|---|---|
| **Yield** | 0.404 (0.100) * | 0.367 (0.105) | 0.404 (0.097) * | 0.383 (0.104) |
| **Height** | 0.500 (0.084) * | 0.426 (0.107) | 0.505 (0.084) * | 0.512 (0.100) * |
| **TKW** | 0.430 (0.096) | 0.423 (0.100) | 0.408 (0.099) | 0.329 (0.099) * |
| **Maturity** | 0.588 (0.066) * | 0.556 (0.080) | 0.547 (0.073) | 0.531 (0.073) * |

TABLE II

Detailed results for the wheat preliminary yield trials. Pearson's r is reported. Standard deviations are shown in brackets. Results significantly higher than GBLUP (one-tailed t-test, p $<0.01$) are denoted by *, and results where GBLUP is significantly higher are denoted by *.

| | GPFN | GBLUP | PCR | XGBoost |
|---|---|---|---|---|
| **Hardness** | 0.681 (0.026) * | 0.667 (0.026) | 0.663 (0.026) | 0.574 (0.029) * |
| **Length** | 0.753 (0.024) * | 0.725 (0.025) | 0.720 (0.024) | 0.671 (0.034) * |
| **PHT** | 0.335 (0.041) | 0.337 (0.040) | 0.326 (0.040) * | 0.233 (0.047) * |
| **PROT** | 0.554 (0.027) * | 0.546 (0.027) | 0.541 (0.027) | 0.453 (0.035) * |
| **TESTW** | 0.630 (0.027) * | 0.612 (0.027) | 0.617 (0.027) | 0.535 (0.031) * |
| **TKW** | 0.669 (0.025) * | 0.648 (0.026) | 0.640 (0.027) * | 0.581 (0.033) * |
| **Width** | 0.736 (0.026) * | 0.717 (0.027) | 0.717 (0.026) | 0.674 (0.026) * |

TABLE III

Detailed results for the wheat landraces. Pearson's r is reported. Standard deviations are shown in brackets. Results significantly higher than GBLUP (one-tailed t-test, p $<0.01$) are denoted by *, and results where GBLUP is significantly higher are denoted by *.

| | GPFN | GBLUP | PCR | XGBoost |
|---|---|---|---|---|
| **DTF** | 0.808 (0.048) * | 0.778 (0.046) | 0.797 (0.045) * | 0.748 (0.053) * |
| **DTM** | 0.755 (0.072) | 0.752 (0.073) | 0.721 (0.079) * | 0.701 (0.084) * |
| **DTS** | 0.819 (0.049) * | 0.774 (0.045) | 0.795 (0.053) * | 0.748 (0.068) * |
| **REP** | 0.600 (0.080) * | 0.562 (0.080) | 0.553 (0.083) | 0.516 (0.091) * |
| **VEG** | 0.817 (0.057) * | 0.784 (0.058) | 0.803 (0.050) * | 0.766 (0.051) * |

TABLE IV

Detailed results for the lentil diversity panel. Pearson's r is reported. Standard deviations are shown in brackets. Results significantly higher than GBLUP (one-tailed t-test, p $<0.01$) are denoted by *, and results where GBLUP is significantly higher are denoted by *.

| | GPFN | GBLUP | PCR | XGBoost |
|---|---|---|---|---|
| **OH Mian** | 0.251 (0.207) | 0.274 (0.208) | 0.215 (0.214) * | 0.259 (0.201) |
| **IL Diers** | 0.115 (0.130) * | 0.080 (0.136) | 0.127 (0.124) * | 0.043 (0.114) * |
| **MI** | -0.112 (0.222) | -0.117 (0.231) | -0.115 (0.206) | -0.146 (0.207) |
| **IA Beavis** | 0.154 (0.138) | 0.141 (0.135) | 0.143 (0.128) | 0.083 (0.121) * |
| **IN** | 0.312 (0.137) | 0.301 (0.137) | 0.289 (0.136) | 0.238 (0.117) * |
| **NE Specht** | 0.301 (0.105) | 0.282 (0.102) | 0.290 (0.102) | 0.187 (0.094) * |
| **OH McHale** | 0.179 (0.097) | 0.174 (0.098) | 0.155 (0.101) | 0.135 (0.089) * |
| **MO Shannon** | 0.226 (0.187) | 0.248 (0.183) | 0.215 (0.198) | 0.169 (0.175) * |
| **KS Schapaugh** | 0.372 (0.132) | 0.355 (0.131) | 0.346 (0.134) | 0.253 (0.125) * |

TABLE V

Detailed results for the between-families soybean NAM population. Pearson's r is reported. Standard deviations are shown in brackets. Results significantly higher than GBLUP (one-tailed t-test, p $<0.01$) are denoted by *, and results where GBLUP is significantly higher are denoted by *.
